# Immunohistochemical Characterization of Phosphorylated Ubiquitin in the Mouse Hippocampus

**DOI:** 10.1101/2020.01.20.912238

**Authors:** Kosuke Kataoka, Andras Bilkei-Gorzo, Andreas Zimmer, Toru Asahi

## Abstract

Mitochondrial autophagy (mitophagy) is an essential and evolutionarily conserved process that maintains mitochondrial integrity via the removal of damaged or superfluous mitochondria in eukaryotic cells. Phosphatase and tensin homolog (PTEN)-induced putative kinase 1 (PINK1) and Parkin promote mitophagy and function in a common signaling pathway. PINK1-mediated ubiquitin phosphorylation at Serine 65 (Ser65-pUb) is a key event in the efficient execution of PINK1/Parkin-dependent mitophagy. However, few studies have used immunohistochemistry to analyze Ser65-pUb in the mouse. Here, we examined the immunohistochemical characteristics of Ser65-pUb in the mouse hippocampus. Some hippocampal cells were Ser65-pUb positive, whereas the remaining cells expressed no or low levels of Ser65-pUb. PINK1 deficiency resulted in a decrease in the density of Ser65-pUb-positive cells, consistent with a previous hypothesis based on *in vitro* research. Interestingly, Ser65-pUb-positive cells were detected in hippocampi lacking PINK1 expression. The CA3 pyramidal cell layer and the dentate gyrus (DG) granule cell layer exhibited significant reductions in the density of Ser65-pUb-positive cells in PINK1-deficient mice. Moreover, Ser65-pUb immunoreactivity colocalized predominantly with neuronal markers. These findings suggest that Ser65-pUb may serve as a biomarker of *in situ* PINK1 signaling in the mouse hippocampus; however, the results should be interpreted with caution, as PINK1 deficiency downregulated Ser65-pUb only partially.

## 1. Introduction

Mitochondrial quality must be surveyed strictly as these organelles regulate numerous biological processes, such as ATP production, lipid metabolism, iron–sulfur cluster biogenesis, and programmed cell death. Mitophagy is an important process that selectively removes damaged or superfluous mitochondria in an autophagic process, to maintain a healthy population of mitochondria and support cell homeostasis. It has become apparent that mitophagy defects can lead to several pathological disorders and that mitophagy-targeting pharmacological or genetical interventions improve such pathologies [1–5].

Recently, significant progress in *in vitro* research has led to the identification of key components of mitophagy, e.g., PINK1 and Parkin, the loss-of-function of which is the most common cause of hereditary early-onset Parkinson’s disease (PD) [6, 7]. These proteins play important roles in the amplification mechanism that increases the efficiency of mitophagy [8]. In healthy mitochondria, after translation in the cytosol, PINK1 is imported into the mitochondrial inner membrane, where it is degraded by mitochondrial-resident proteases [9]. However, mitochondrial damage stemming from mutations in mitochondrial DNA and the accumulation of misfolded proteins [10, 11] leads to the stabilization of PINK1 in the outer membrane [12–14]. Importantly, activated PINK1 phosphorylates (at Serine 65) the ubiquitin molecules that are attached to proteins in the outer membrane [15–17]. The resulting Ser65-pUb recruits the E3 ubiquitin ligase Parkin from the cytosol to mitochondria, where Parkin is phosphorylated by PINK1 to potentiate its latent E3 ligase activity [18–21]. Finally, a positive feedback loop accelerates Parkin translocation and the formation of poly-ubiquitin chains on damaged mitochondria. Amplified Ser65-pUb chains recruit various factors that are required for autophagy, to execute mitophagy [8].

Although the precise mechanisms underlying the PINK1/Parkin-dependent mitophagy pathway have been intensively investigated in *in vitro* studies, many questions remain pertaining to the existence and physiological relevance of mitophagy *in vivo*. In *Drosophila*, mitophagy is present in muscles and brain tissues in physiological conditions [22]. *Drosophila* lacking *PINK1* or *Parkin* exhibit mitochondrial, neurological, and behavioral abnormalities that closely resemble the symptoms of human patients with PD [23–26]. Ser65-pUb expression is also observed in dopaminergic neurons in *Drosophila*, suggesting that PINK1/Parkin signaling functions faithfully [27]. A study using human postmortem brain specimens showed that Ser65-pUb is widely distributed throughout the brain, including the substantia nigra and hippocampus, and that Ser65-pUb is upregulated in an age-dependent manner and in PD brains [28, 29]. In addition, Ser65-pUb generation is impaired in iPSCs derived from PD patients harboring *PINK1* and *Parkin* mutations [27, 29]. In contrast to fly models, *PINK1* and *Parkin* knockout mice failed to exhibit PD pathologies, despite the presence of pronounced mitochondrial dysfunction (reviewed in [30]). Moreover, a reporter mouse model for *in vivo* mitophagy showed that basal mitophagy occurs independently of PINK1 [31]. Nevertheless, a recent study using an elegant genetic mouse model highlighted the importance of endogenous mitophagy activity in the mouse [32], as a synergistic effect of Parkin deficiency and genetic defects in mitochondrial DNA proofreading leads to the accumulation of Ser65-pUb in brains, as measured using quantitative mass spectrometry, which does not provide spatial information [32]. Although the importance of PINK1/Parkin signaling at the basal level has been re-emphasized in mouse models, little is known about the *in situ* PINK1/Parkin signaling activity in mouse tissues.

Here, we investigated immunohistochemically Ser65-pUb in mouse hippocampi, because Ser65-pUb is emerging as a possible biomarker of PINK1 signaling activity *in vivo* [27, 29, 33]. Our data provide insights into the physiological role of Ser65-pUb in the mouse and suggest that Ser65-pUb is useful for monitoring *in situ* PINK1 signaling activity in mouse tissues.

## 2. Materials and Methods

### 2.1. Animals

Brains obtained from *PINK1*^−/−^ and *PINK1*^+/+^ mice (aged 7 months) were kindly gifted by Dr. Miratul Muqit (MRC Protein Phosphorylation Unit, University of Dundee) and stored at −80°C until further processing. The animals used in all experiments were mix-gendered. *PINK1*^−/−^ mice were generated as described previously [34].

### 2.2. Tissue preparation

Brain samples containing the hippocampus were cut serially into 16-μm-thick coronal sections using a cryostat (Leica CM 3050; Leica, Welzlar, Germany) and mounted onto glass slides. Slides were stored at −80°C until staining.

### 2.3. Immunofluorescence staining

Frozen sections were placed on a hot plate at 40°C for 20–30 min. Brain sections were framed with a PapPen and post-fixed in 4% paraformaldehyde dissolved in 0.1 M phosphate-buffered saline (PBS; pH 7.3) for 30 min. After three 10-min washes in PBS, sections were permeabilized in 0.5% Triton X-100 for 1 h, followed by three 10-min washes in PBS. Blocking of unspecific binding sites was performed by incubation in PBS containing 3% bovine serum albumin (BSA) for 2 h. The slices were then incubated with a rabbit anti-phospho-ubiquitin (Ser65) antibody (Merck Millipore, Burlington, MA, USA, ABS1513-l; dilution, 1:500 in 0.3% BSA in PBS) and an Alexa Fluor 488-conjugated mouse anti-NeuN antibody (Merck Millipore, MAB377X; dilution, 1:500 in 0.3% BSA in PBS) at 4°C overnight. Subsequently, the slides were rinsed three times for 10 min in PBS and incubated with a goat anti-rabbit Cy3-conjugated secondary antibody (Invitrogen, Waltham, CA, USA, 1081937; dilution, 1:1000 in 0.3% BSA in PBS). After staining, the sections were rinsed three times for 10 min in PBS, mounted with DAPI-containing Fluoromount-G (Southern Biotech, Birmingham, AL, USA), and covered with cover slips.

### 2.4. Confocal imaging

To assess the colocalization of Ser65-pUb/NeuN and Ser65-pUb/lipofuscin, images were acquired using a Leica LSM SP8 confocal microscope equipped with a 60× water-immersion lens. For the analysis of Ser65-pUb-positive cells in the whole hippocampus, images were acquired using a Leica LSM SP8 confocal microscope equipped with a 20× lens in the “Tile Scan” mode. Three to four images per animal that included the whole hippocampus were obtained for analysis. The number of Ser65-pUb-positive cells in the hippocampus was counted automatically using the “Analyze particles” tool in Image J Fiji (NIH, Bethesda, MD, USA), after thresholding. For the analysis of Ser65-pUb and DAPI signal intensity in the CA3 region, images were acquired using a Leica LSM SP8 confocal microscope equipped with a 10× lens. Three to four images per sample that included the whole CA3 region were obtained for analysis. Finally, the CA3 regions in the hippocampus were framed using the “Polygon selection” tool and Ser65-pUb signal intensities were measured using Image J Fiji (NIH).

### 2.5. Statistics

Statistical analysis was performed using GraphPad Prism (version 5.0 or 8.0). To analyze the number of Ser65-pUb-positive cells in the whole hippocampus, data were processed using unpaired Student’s *t*-tests with Welch’s correction, because they showed significantly unequal variances. The remaining data were analyzed using unpaired Student’s *t*-tests. \**P* < 0.05, \*\**P* < 0.01, significant differences compared with the *PINK1*^+/+^ counterparts.

## 3. Results

First, we investigated the immunohistochemical characteristics of Ser65-pUb in the mouse hippocampus. Some cells exhibiting strong signals were distributed throughout the hippocampus, whereas the remainder of the cells expressed no or weak Ser65-pUb in *PINK1*^+/+^ mice (Fig. 1A). We used a genetic mouse model to confirm that these immunoreactivities were the consequence of specific staining for Ser65-pUb [34]. As PINK1 catalyzes the phosphorylation of ubiquitin at Serine 65 for the execution of mitophagy [15–17], next we checked whether PINK1 deficiency affected Ser65-pUb staining in the hippocampus. To avoid analytical inaccuracies, we performed unbiased image quantification, i.e., Ser65-pUb-positive cells in the hippocampus were counted after the application of image segmentation with the same threshold range across all images. This method revealed that PINK1 deficiency resulted in a significant decrease in the density of Ser65-pUb-positive cells in the whole hippocampal area (Fig. 1A and B), indicating that Ser65-pUb positivity in these cells resulted, at least partly, from PINK1 activity in the hippocampus. Of note, Ser65-pUb-positive cells were also detected in *PINK1*^−/−^ mice. To confirm that PINK1 contributes to the formation of Ser65-pUb in the hippocampus, we also measured Ser65-pUb fluorescence intensities in the hippocampal CA3 area. The Ser65-pUb signal intensity in the CA3 area was significantly decreased in *PINK1*^−/−^ mice compared with age-matched *PINK1*^+/+^ mice, whereas the DAPI signal was comparable between mice of the two genotypes (Supplementary Fig. 1). This indicates that, in the hippocampus of *PINK1*^−/−^ mice, Ser65-pUb expression was decreased while the cell density was unchanged compared with the *PINK1*^+/+^ counterparts. Overall, these findings suggest that Ser65-pUb is produced at high levels in a subset of cells of the hippocampus, which is at least partially dependent on PINK1.

**Fig. 1.**
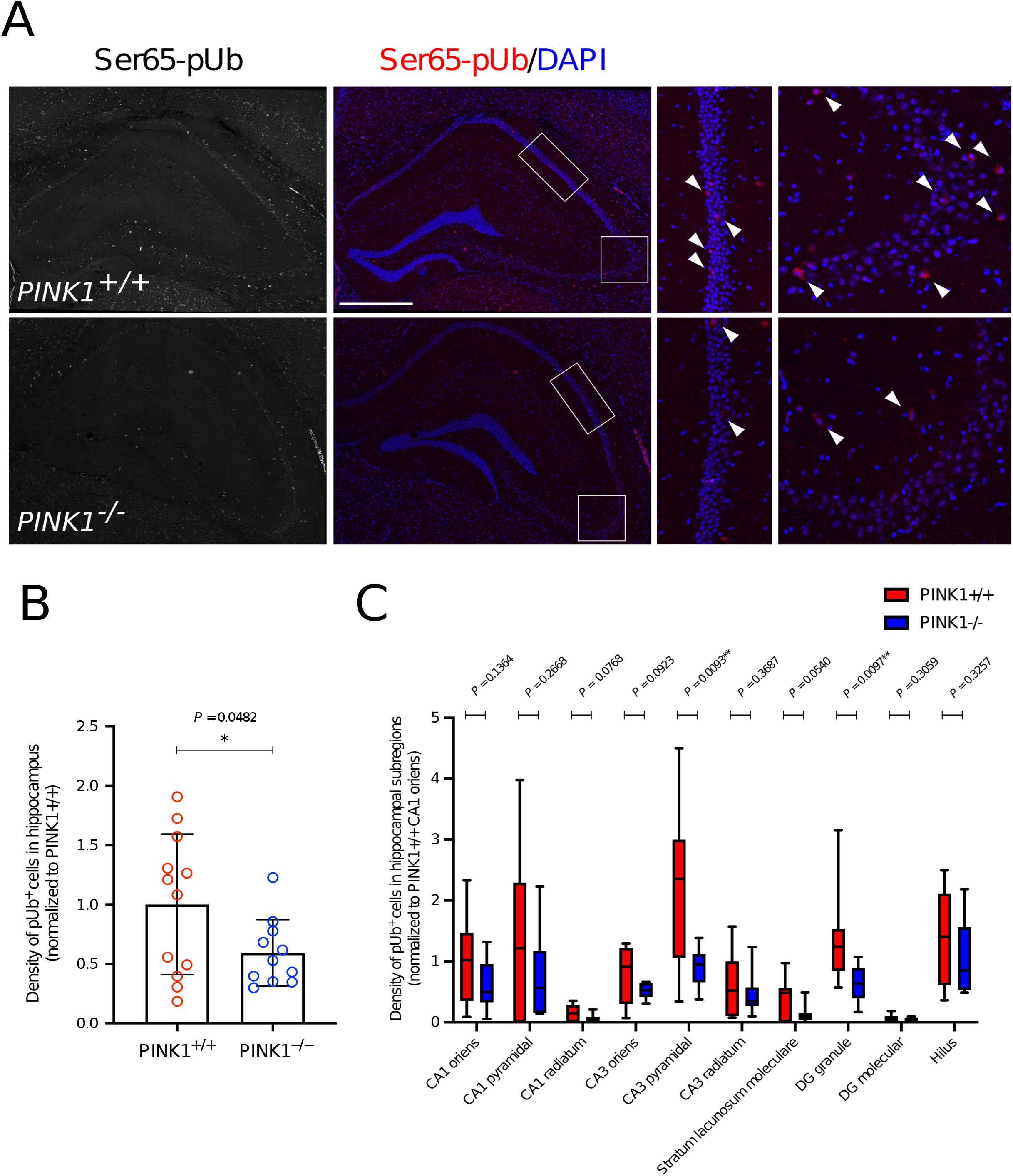
Ser65-pUb-positive cells in the mouse hippocampus. (A) Representative confocal microscopy images of hippocampi in *PINK1*^+/+^ and *PINK1*^−/−^ mice. Coronal sections were immunostained with an anti-Ser65-pUb antibody (red). Nuclei were stained with DAPI (blue). The boxed regions are magnified on the right. The white arrows indicate examples of cells strongly expressing Ser65-pUb. Scale bar, 500 μm. (B) Quantitative analysis of the density of Ser65-pUb-positive cells in the whole hippocampus in *PINK1*^+/+^ and *PINK1*^−/−^ mice. The results are expressed as the mean ± S.E.M. The data points represent the density of Ser65-pUb-positive cells in one section. Three animals per genotype were used in this analysis. (C) Quantitative analysis of the density of Ser65-pUb-positive cells in the indicated hippocampal subregions. The box plots represent minimum to maximum values, with the box denoting the 25th, 50th (median), and 75th percentile.

The hippocampal body is composed of the CA and DG regions. These regions can be further divided into 10 subregions that are anatomically distinct (Supplementary Fig. 2A). These hippocampal subregions play important cooperative or uncooperative roles in memory and learning. To examine the subregion in which Ser65-pUb expression is affected by PINK1 deficiency, next we measured the density of Ser65-pUb-positive cells in each hippocampal subregion in *PINK1*^+/+^ and *PINK1*^−/−^ mice. In *PINK1*^+/+^ mice, Ser65-pUb-positive cells were observed particularly in the CA1/3 oriens and pyramidal cell layers, DG granule cell layer, and hilus and, to a lesser extent, in the CA3 radiatum and stratum lacunosum moleculare (Fig. 1C). The CA1 radiatum and DG molecular layer exhibited almost no Ser65-pUb-positive cells (Fig. 1C). In *PINK1*^−/−^ mice, the density of Ser65-pUb-positive cells was significantly reduced especially in the CA3 pyramidal cell layer and DG granule cell layer (Fig. 1C). PINK1 deficiency did not have an effect on the proportion of Ser65-pUb-positive cells in each area (Supplementary Fig. 2B). These data indicate that *PINK1* knockout affects Ser65-pUb expression in specific hippocampal subregions.

As the animals used in this experiment were approximately 7 months old, lipofuscin autofluorescence might affect our interpretation of Ser65-pUb immunoreactivity. However, we could distinguish lipofuscin autofluorescence from Ser65-pUb fluorescence (Supplementary Fig. 3). We also noticed that the nuclei of cells in the CA1/3 pyramidal cell layers were immunoreactive against Ser65-pUb in both *PINK1*^+/+^ and *PINK1*^−/−^ mice. (Supplementary Fig. 4A). Nevertheless, we found no differences between the two genotypes regarding Ser65-pUb signal intensities in DAPI-positive areas in pyramidal cell layers (Supplementary Fig. 4B), indicating that the immunoreactivity observed in nuclei in CA1/3 pyramidal cell layers might be unspecific.

As the brain comprises neurons and glial cells, such as astrocytes, next we examined the type of hippocampal cells that express Ser65-pUb. Ser65-pUb was expressed predominantly in NeuN-positive cells (neurons), and not in GFAP-positive cells (astrocytes), in the hippocampus (Fig. 2A and B). These results indicate that, in physiological conditions, hippocampal neurons express Ser65-pUb in a PINK1-dependent manner.

**Fig. 2.**
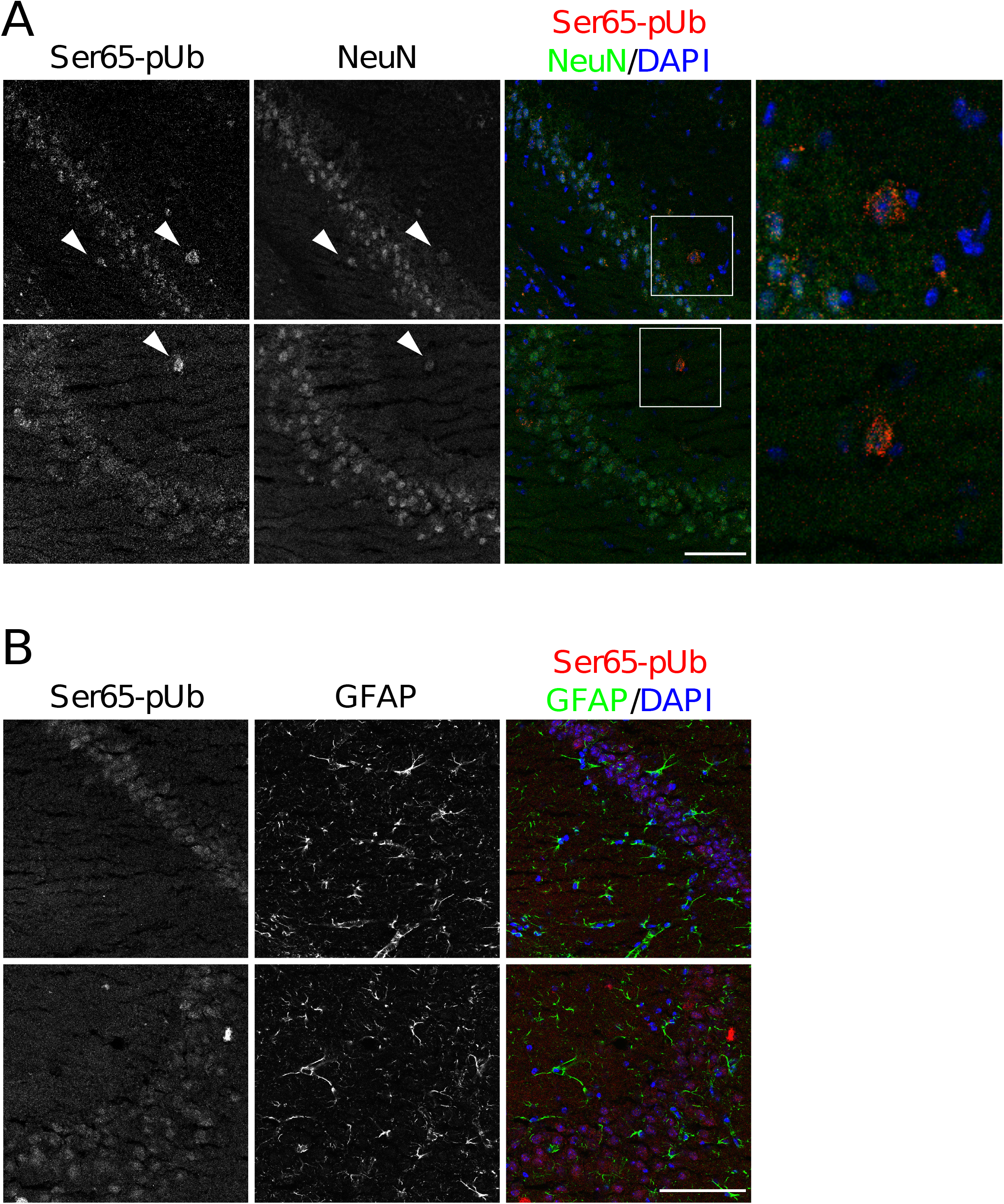
Colocalization of Ser65-positive cells with neuronal and astrocytic markers. (A) Representative confocal microscopy images of hippocampal CA1 (upper images) and CA3 (lower images) regions. Coronal sections were immunostained with anti-Ser65-pUb (red), and anti-NeuN (green) antibodies. Nuclei were stained with DAPI (blue). The boxed regions are magnified on the right. The white arrows indicate Ser65-pUb-positive cells expressing NeuN. Scale bar, 100 μm. (B) Representative confocal microscopy images of hippocampal CA1 (upper images) and CA3 (lower images) regions. Coronal sections were immunostained with anti-Ser65-pUb (red) and anti-GAFP (green) antibodies. Nuclei were stained with DAPI (blue). Scale bar, 100 μm.

## 4. Discussion

Ubiquitin phosphorylation at Serine 65 is a key step in the efficient execution of PINK1/Parkin-dependent mitophagy [8]. The physiological and pathological importance of ubiquitin phosphorylation has been suggested [27–29]. However, immunohistochemical data on ubiquitin phosphorylation are scarce in mouse models. We investigated the immunohistochemical characteristics of Ser65-pUb in the mouse hippocampus. Some cells expressed Ser65-pUb strongly throughout the hippocampus, while other cells expressed no or low levels of Ser65-pUb. These Ser65-pUb-positive cells were significantly decreased in *PINK1* knockout mice, indicating that PINK1 might be physiologically active in the hippocampus. Moreover, the CA3 pyramidal cell layer and DG granule cell layer were significantly affected by PINK1 deficiency. Interestingly, a morphometric analysis of the subcortical gray structure in PD patients showed that the CA3 and DG, rather than the CA1, exhibit atrophy [35, 36]. This raises the possibility that PINK1-mediated Ser65-pUb formation is important for a proper structure and function of the hippocampus in the context of PD. Moreover, Ser65-pUb was mainly expressed in neurons. Consistently, *in situ* hybridization revealed that the *PINK1* mRNA is strongly expressed in neurons, especially in the hippocampal CA3 layer, with little to no expression in glial cells in the rodent brain [37, 38]. Notably, mitophagy is important for proper neuronal function and survival via the maintenance of a healthy pool of mitochondria [39, 40]. Thus, Ser65-pUb might play an important role in preserving neurons in the mouse hippocampus through mitophagy.

PINK1 is the only known kinase to phosphorylate ubiquitin at Serine 65. Although we showed that PINK1 deficiency resulted in a significant reduction in the density of Ser65-pUb-positve cells, several Ser65-pUb-positive cells were detected in the hippocampus of *PINK1*^−/−^ mice. Notably, PINK1-independent Ser65-pUb signals were observed in regions adjacent to the PPM2 cluster in *Drosophila* lacking *PINK1* [27]. These findings suggest that one or more uncharacterized kinases other than PINK1 produce Ser65-pUb. However, we cannot exclude the possibility that this PINK1-independent fluorescence signal results from unspecific staining.

The decreased density of Ser65-pUn-positive cells may not result from PINK1 deficiency; rather, it may be attributed to a reduction in neuronal density. Previously, loss of PINK1 was shown to impede the differentiation of neuronal stem cells through mitochondrial defects in the DG subgranular zone of the hippocampus [41]. However, the same study demonstrated that neuronal density in the hippocampus is unchanged in *PINK1* knockout mice [41]. Moreover, several studies have suggested that *PINK1*^−/−^ mice do not exhibit major neurodegeneration, despite the presence of pronounced mitochondrial dysfunction (reviewed in [30]). These studies support the attribution of the reduction in Ser65-pUb-positive cell density in PINK1 knockout mice to a loss of PINK1 activity in neurons, rather than a reduction in neuronal density.

We found that a subset of the cells strongly express Ser65-pUb while other cells do not, and that PINK1 is responsible for Ser65-pUb formation in the mouse, similar to that observed in a fly model and human subjects [27–29]. The neuronal expression of Ser65-pUb suggests an important role for this protein in maintaining neuronal mitochondrial integrity. Our findings showed that Ser65-pUb may be useful for biomarker of *in situ* PINK1 activity in the mouse hippocampus. However, these results should be interpreted cautiously, as a substantial number of Ser65-pUb-positive cells was detected in mouse hippocampi lacking PINK1, indicating that uncharacterized kinase(s) might regulate the expression of Ser65-pUb. Given that PINK1 expression is prevalent not only in the brain, but also in peripheral tissues (such as the heart), further analyses of Ser65-pUb in different organs of the mouse are warranted.

## Acknowledgments

We thank Dr. Chihiro Nozaki for technical supports and fruitful discussions; members of Zimmer and Asahi laboratory for suggestions. We also thank Professor Miratul Muqit, University of Dundee, for kindly giving us PINK1 brains. This study was supported by the Grant-in-Aid for Early-Career Scientists (19K20196, K.K.), the Top Global University Project from MEXT, Japan and Tokutei Kadai, Newly Hired Faculty from Waseda university (BARA001611, K.K.).

## Author Contributions

K. K. and A. B. G. designed experiments. K. K. performed all experiments, analyzed data, and wrote the manuscript. A. B. G., A. Z., and T. A. conceived and supervised the project. All authors read and approved the final manuscript.

## Notes

The authors declare that they have no conflicts of interest.

## References

[1] A. Rana, M. Rera, D. W. Walker, Parkin overexpression during aging reduces proteotoxicity, alters mitochondrial dynamics, and extends lifespan, Proc. Natl. Acad. Sci. U.S.A. 110 (2013) 8638–8643. https://doi.org/10.1073/pnas.1216197110.

[2] E. F. Fang, Y. Hou, K. Palikaras et al. Mitophagy inhibits amyloid-β and tau pathology and reverses cognitive deficits in models of Alzheimer’s disease, Nat. Neurosci. 22 (2019) 401–412. https://doi.org/10.1038/s41593-018-0332-9.

[3] V. Sorrentino, M. Romani, L. Mouchiroud et al. Enhancing mitochondrial proteostasis reduces amyloid-β proteotoxicity, Nature 552 (2017) 187–193. https://doi.org/10.1038/nature25143.

[4] A. Hoshino, Y. Mita, Y. Okawa et al. Cytosolic p53 inhibits Parkin-mediated mitophagy and promotes mitochondrial dysfunction in the mouse heart, Nat. Commun. 4 (2013) 1–12. https://doi.org/10.1038/ncomms3308.

[5] D. Ryu, L. Mouchiroud, P. A. Andreux et al. Urolithin A induces mitophagy and prolongs lifespan in C. elegans and increases muscle function in rodents, Nat. Med. 22 (2016) 879–888. https://doi.org/10.1038/nm.4132.

[6] T. Kitada, S. Asakawa, N. Hattori et al. Mutations in the parkin gene causeautosomal recessive juvenile parkinsonism, Nature 169 (1998) 166–169. https://doi.org/10.1038/33416.

[7] E. M. Valente, P. M. Abou-Sleiman, V. Caputo et al. Hereditary early-onset Parkinson’s Disease caused by mutations in PINK1, Science 304 (2004) 1158–1160. https://doi.org/10.1126/science.1096284.

[8] M. Lazarou, D. A. Sliter, L. A. Kane et al. The ubiquitin kinase PINK1 recruits autophagy receptors to induce mitophagy, Nature 524 (2015) 309–314. https://doi.org/10.1038/nature14893.

[9] R. E. Thomas, L. A. Andrews, J. L. Burman et al. PINK1-Parkin Pathway activity is regulated by degradation of PINK1 in the mitochondrial matrix, PLoS Genet. 10 (2014) e1004279. https://doi.org/10.1371/journal.pgen.1004279.

[10] R. R. H. Hämäläinen, T. Manninen, H. Koivumäki et al. Tissue-and cell-type–specific manifestations of heteroplasmic mtDNA 3243A>G mutation in human induced pluripotent stem cell-derived disease model, Proc. Natl. Acad. Sci. U.S.A. 110 (2013) 3622–3630. https://doi.org/10.1073/pnas.1311660110.

[11] J. L. Burman, S. Pickles, C. Wang et al. Mitochondrial fission facilitates the selective mitophagy of protein aggregates, J. Cell Biol. 216 (2017) 3231–3247. https://doi.org/10.1083/jcb.201612106.

[12] N. Matsuda, S. Sato, K. Shiba et al. PINK1 stabilized by mitochondrial depolarization recruits Parkin to damaged mitochondria and activates latent Parkin for mitophagy, J. Cell Biol. 189 (2010) 211–221. https://doi.org/10.1083/jcb.200910140.

[13] D. P. Narendra, S. M. Jin, A. Tanaka et al. PINK1 is selectively stabilized on impaired mitochondria to activate Parkin, PLoS Biol. 8 (2010) e1000298. https://doi.org/10.1371/journal.pbio.1000298.

[14] S. M. Jin, M. Lazarou, C. Wang et al. Mitochondrial membrane potential regulates PINK1 import and proteolytic destabilization by PARL, J. Cell Biol. 191 (2010) 933–942. https://doi.org/10.1083/jcb.201008084.

[15] F. Koyano, K. Okatsu, H. Kosako et al. Ubiquitin is phosphorylated by PINK1 to activate parkin, Nature 510 (2014) 162–166. https://doi.org/10.1038/nature13392.

[16] L. A. Kane, M. Lazarou, A. I. Fogel et al. PINK1 phosphorylates ubiquitin to activate Parkin E3 ubiquitin ligase activity, J. Cell Biol. 205 (2014) 143–153. https://doi.org/10.1083/jcb.201402104.

[17] A. Kazlauskaite, C. Kondapalli, R. Gourlay et al. Parkin is activated by PINK1-dependent phosphorylation of ubiquitin at Ser 65, Biochem. J. 139 (2014) 127–139. https://doi.org/10.1042/BJ20140334.

[18] M. Iguchi, Y. Kujuro, K. Okatsu et al. Parkin-catalyzed ubiquitin-ester transfer is triggered by PINK1-dependent phosphorylation, J. Biol. Chem. 288 (2013) 22019–22032. https://doi.org/10.1074/jbc.M113.467530.

[19] K. Shiba-fukushima, Y. Imai, S. Yoshida et al. PINK1-mediated phosphorylation of the Parkin ubiquitin-like domain primes mitochondrial translocation of Parkin and regulates mitophagy, Sci. Rep. (2012) 1002. https://doi.org/10.1038/srep01002.

[20] C. Kondapalli, A. Kazlauskaite, N. Zhang et al. PINK1 is activated by mitochondrial membrane potential depolarization and stimulates Parkin E3 ligase activity by phosphorylating Serine 65, Open Biol. 2 (2012) 120080. https://doi.org/10.1098/rsob.120080.

[21] T. G. Mcwilliams, M. M. K. Muqit, E. Barini et al. Phosphorylation of Parkin at serine 65 is essential for its activation in vivo, Open Biol. 8 (2018) 180108. https://doi.org/10.1098/rsob.180108

[22] T. Cornelissen, S. Vilain, K. Vints et al. Deficiency of parkin and PINK1 impairs age-dependent mitophagy in Drosophila, Elife 7 (2018) e35878. https://doi.org/10.7554/eLife.35878.

[23] I. E. Clark, M. W. Dodson, C. Jiang et al. Drosophila pink1 is required for mitochondrial function and interacts genetically with parkin, Nature 441 (2006) 1162–1166. https://doi.org/10.1038/nature04779.

[24] J. C. Greene, A. J. Whitworth, I. Kuo et al. Mitochondrial pathology and apoptotic muscle degeneration in Drosophila parkin mutants, Proc. Natl. Acad. Sci. U.S.A. 100 (2003) 4078–4083. https://doi.org/10.1073/pnas.0737556100.

[25] J. Park, S. B. Lee, S. Lee et al. Mitochondrial dysfunction in Drosophila PINK1 mutants is complemented by parkin, Nature 441 (2006) 1157–1161. https://doi.org/10.1038/nature04788.

[26] A. J. Whitworth, D. A. Theodore, J. C. Greene et al. Increased glutathione S-transferase activity rescues dopaminergic neuron loss in a Drosophila model of Parkinson’s disease, Proc. Natl. Acad. Sci. U.S.A. 102 (2005) 8024–8029. https://doi.org/10.1073/pnas.0501078102.

[27] K. Shiba-Fukushima, K. I. Ishikawa, T. Inoshita et al. Evidence that phosphorylated ubiquitin signaling is involved in the etiology of Parkinson’s disease, Hum. Mol. Genet. 26 (2017) 3172–3185. https://doi.org/10.1093/hmg/ddx201.

[28] F. C. Fiesel, M. Ando, R. Hudec et al. (Patho-)physiological relevance of PINK 1-dependent ubiquitin phosphorylation, EMBO Rep. 16 (2015) 1114–1130. https://doi.org/10.15252/embr.201540514.

[29] X. Hou, F. C. Fiesel, D. Truban et al. Age- and disease-dependent increase of the mitophagy marker phospho-ubiquitin in normal aging and Lewy body disease, Autophagy 14 (2018) 1404–1418. https://doi.org/10.1080/15548627.2018.1461294.

[30] T. M. Dawson, H. S. Ko, V. L. Dawson, Genetic animal models of Parkinson’s Disease, Neuron 66 (2010) 646–661. https://doi.org/10.1016/j.neuron.2010.04.034.

[31] T. G. McWilliams, A. R. Prescott, L. Montava-Garriga et al. Basal mitophagy occurs independently of PINK1 in Mouse tissues of high metabolic demand, Cell Metab. 27 (2018) 439–449. https://doi.org/10.1016/j.cmet.2017.12.008.

[32] A. M. Pickrell, C. H. Huang, S. R. Kennedy et al. Endogenous Parkin Preserves dopaminergic substantia nigral neurons following mitochondrial DNA mutagenic stress, Neuron 87 (2015) 371–382. https://doi.org/10.1016/j.neuron.2015.06.034.

[33] D. A. Sliter, J. Martinez, L. Hao et al. Parkin and PINK1 mitigate STING-induced inflammation, Nature 561 (2018) 258–262. https://doi.org/10.1038/s41586-018-0448-9.

[34] A. Wood-kaczmar, S. Gandhi, Z. Yao et al. PINK1 Is necessary for long term survival and mitochondrial function in human dopaminergic neurons, PLoS One 3 (2008) e2455. https://doi.org/10.1371/journal.pone.0002455.

[35] J. B. Pereira, C. Junqu, D. Bartr et al. Regional vulnerability of hippocampal subfields and memory deficits in Parkinson’s Disease, Hippocampus 728 (2013) 720–728. https://doi.org/10.1002/hipo.22131.

[36] J. J. Tanner, N. R. Mcfarland, C. C. Price et al. Striatal and hippocampal atrophy in idiopathic Parkinson’s Disease patients without dementia: A morphometric analysis, Front. Neurol. 8, (2017) https://doi.org/10.3389/fneur.2017.00139.

[37] J. M. Taymans, C. Van Den Haute, V. Baekelandt, Distribution of PINK1 and LRRK2 in rat and mouse brain, J. Neurochem. 98 (2006) 951–961. https://doi.org/10.1111/j.1471-4159.2006.03919.x.

[38] J. G. Blackinton, A. Anvret, A. Beilina et al. Expression of PINK1 mRNA in human and rodent brain and in Parkinson’s disease, Brain Res. 1184 (2007) 10–16. https://doi.org/10.1016/j.brainres.2007.09.056.

[39] E. F. Fang, H. Kassahun, D. L. Croteau et al. NAD+ replenishment improves lifespan and healthspan in ataxia telangiectasia models via mitophagy and DNA Repair, Cell Metab. 24 (2016) 566–581. https://doi.org/10.1016/j.cmet.2016.09.004.

[40] E. F. Fang, M. Scheibye-knudsen, L. E. Brace et al. Defective mitophagy in XPA via PARP-1 hyperactivation and NAD+/SIRT1 reduction, Cell 157 (2014) 882–896. https://doi.org/10.1016/j.cell.2014.03.026.

[41] S. K. Agnihotri, R. Shen, J. Li et al. Loss of PINK1 leads to metabolic deficits in adult neural stem cells and impedes differentiation of newborn neurons in the mouse hippocampus, FASEB J. 31 (2017) 2839–2853. https://doi.org/10.1096/fj.201600960RR.

