## Supplemental Figures for "Immunohistochemical Characterization of Phosphorylated Ubiquitin in the Mouse Hippocampus"

### Supplemental Fig. 1

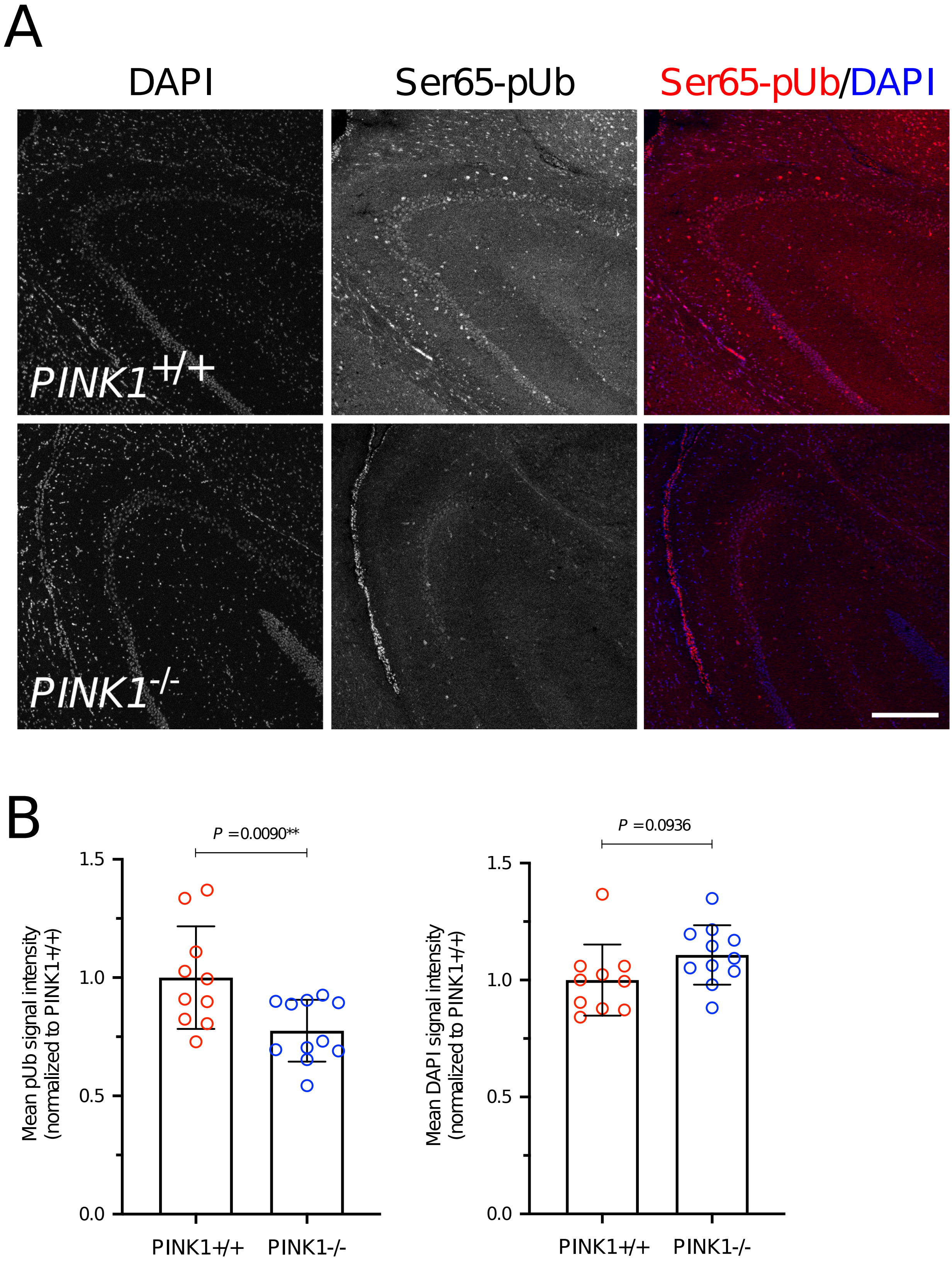

**Supplementary Figure 1. CA3 regions in *PINK1*$^{+/+}$ and *PINK1*$^{-/-}$ mouse** (A) Representative confocal microscopy images of hippocampal CA3 regions in PINK1$^{+/+}$ and PINK1$^{-/-}$ mouse. Coronal sections were immunostained with Ser65-pUb antibody (Red). Nucleus was stained with DAPI (Blue). Scale bar: 250 μm. (B) Quantitative analysis of fluorescence signal intensities of Ser65-pUb and DAPI in CA3 area between PINK1$^{+/+}$ and PINK1$^{-/-}$. Data points represent mean fluorescence signal intensity in one section. Three animals per genotype were used for the analysis.

### Supplemental Fig. 2

**A**

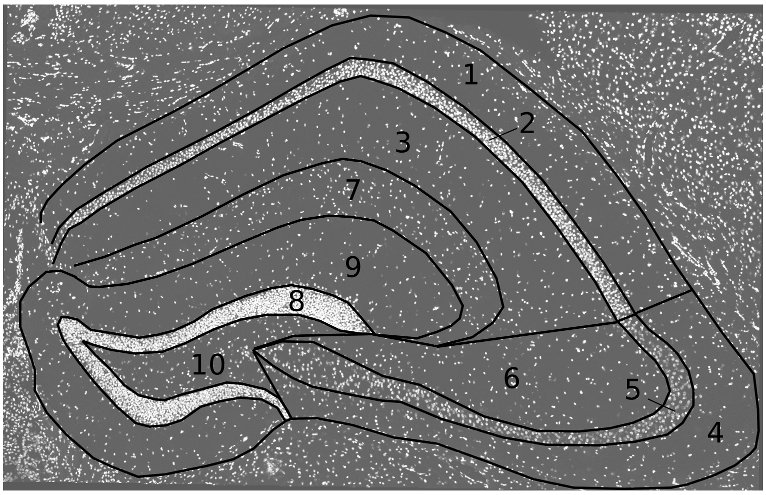


1. CA1 oriens
2. CA1 pyramidal
3. CA1 radiatum
4. CA3 oriens
5. CA3 pyramidal
6. CA3 radiatum
7. Stratum lacunosum moleculare
8. DG granule
9. DG molecular
10. Hilus

**B**

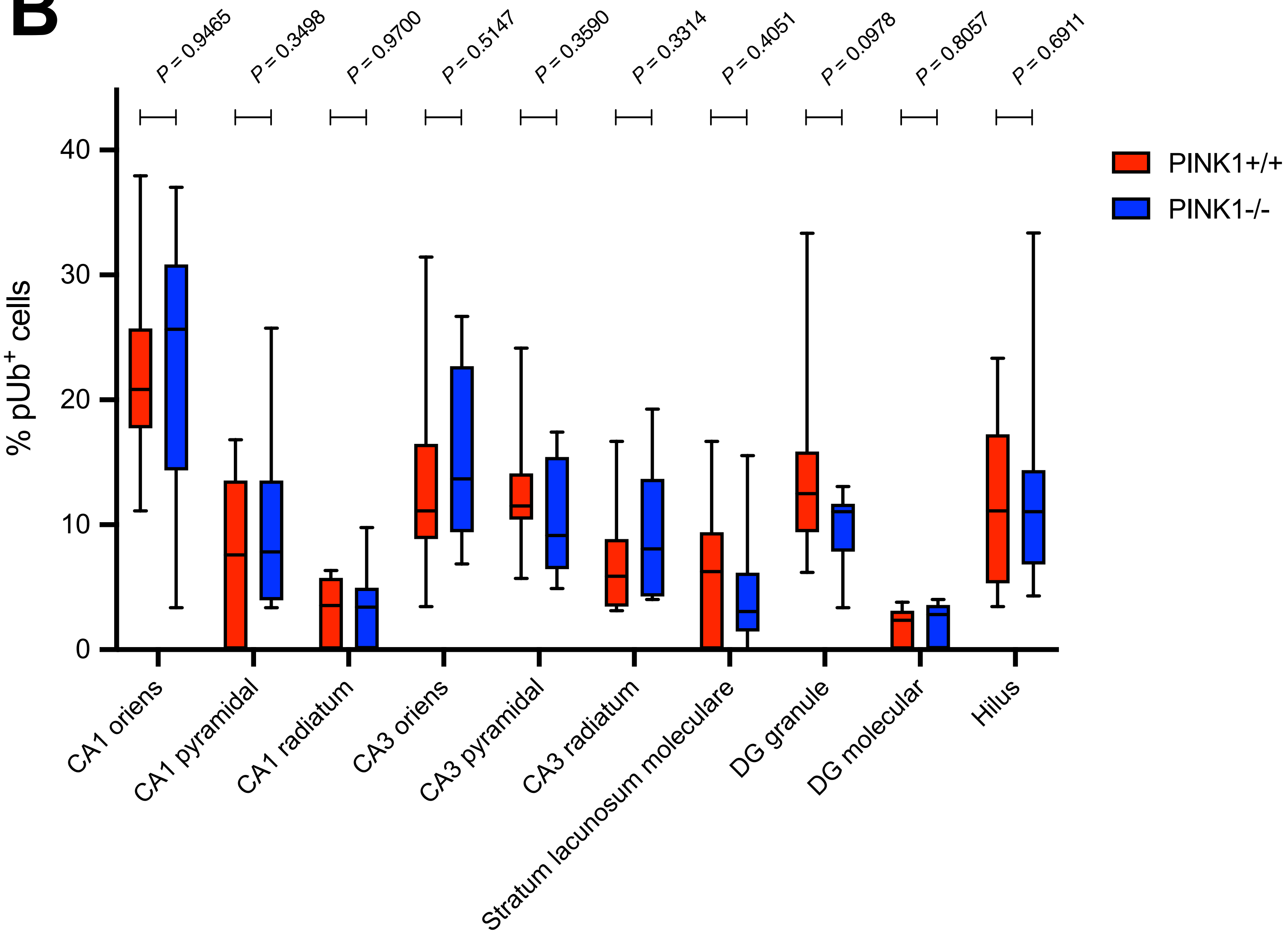

**Supplementary Figure 2. Subregional analysis of Ser65-pUb-positive cells in hippocampus** (A) Ten subregions in hippocampus are shown based on anatomical characteristics from DAPI and Ser65-pUb images. (B) Quantitative analysis of proportion of Ser65-pUb-positive cells in indicated hippocampal subregions. Box plots represent minimum to maximum values, with the box denoting 25th, 50th (median) and 75th percentile.

### Supplemental Fig. 3

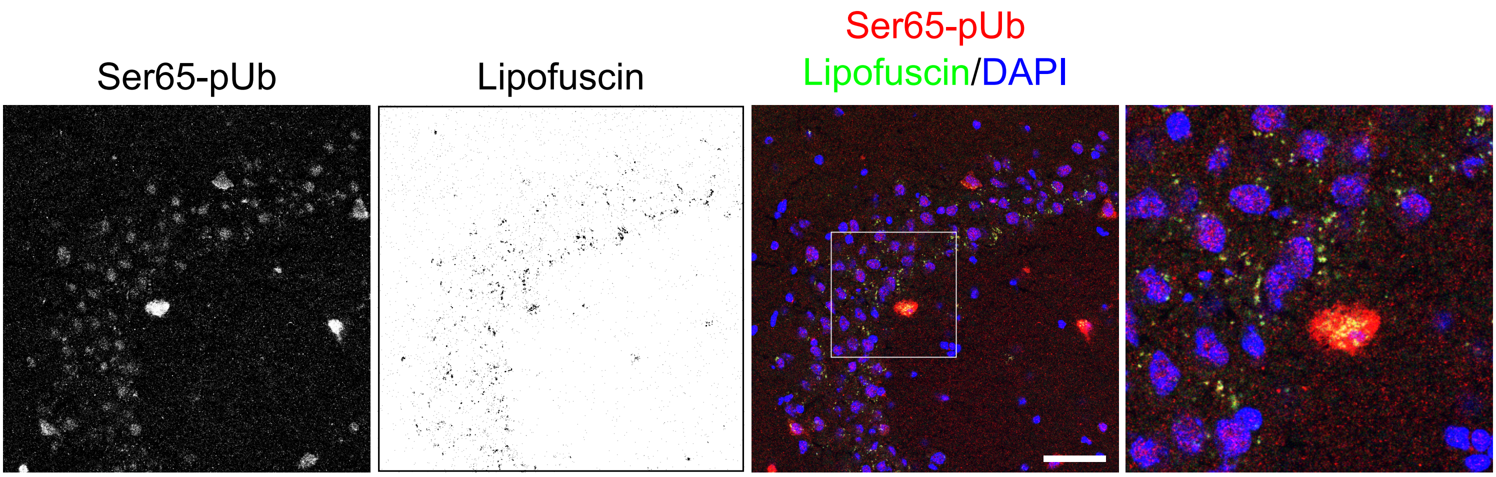

**Supplementary Figure 3. Localization of lipofuscin autofluorescence and Ser65-pUb signal** Representative confocal microscopy images of hippocampal CA3 region. Coronal sections were immunostained with anti-Ser65-pUb (Red). Lipofuscin autofluorescence signal was obtained by acquiring the overlapping pixels of emission at 510-530 nm (Green) and at 570-600 nm (Red) with excitation at 488 nm and 561 nm, respectively. Nucleus was stained with DAPI (Blue). Boxed regions are shown magnified on the right. Scale bar: 20 μm.

### Supplemental Fig. 4

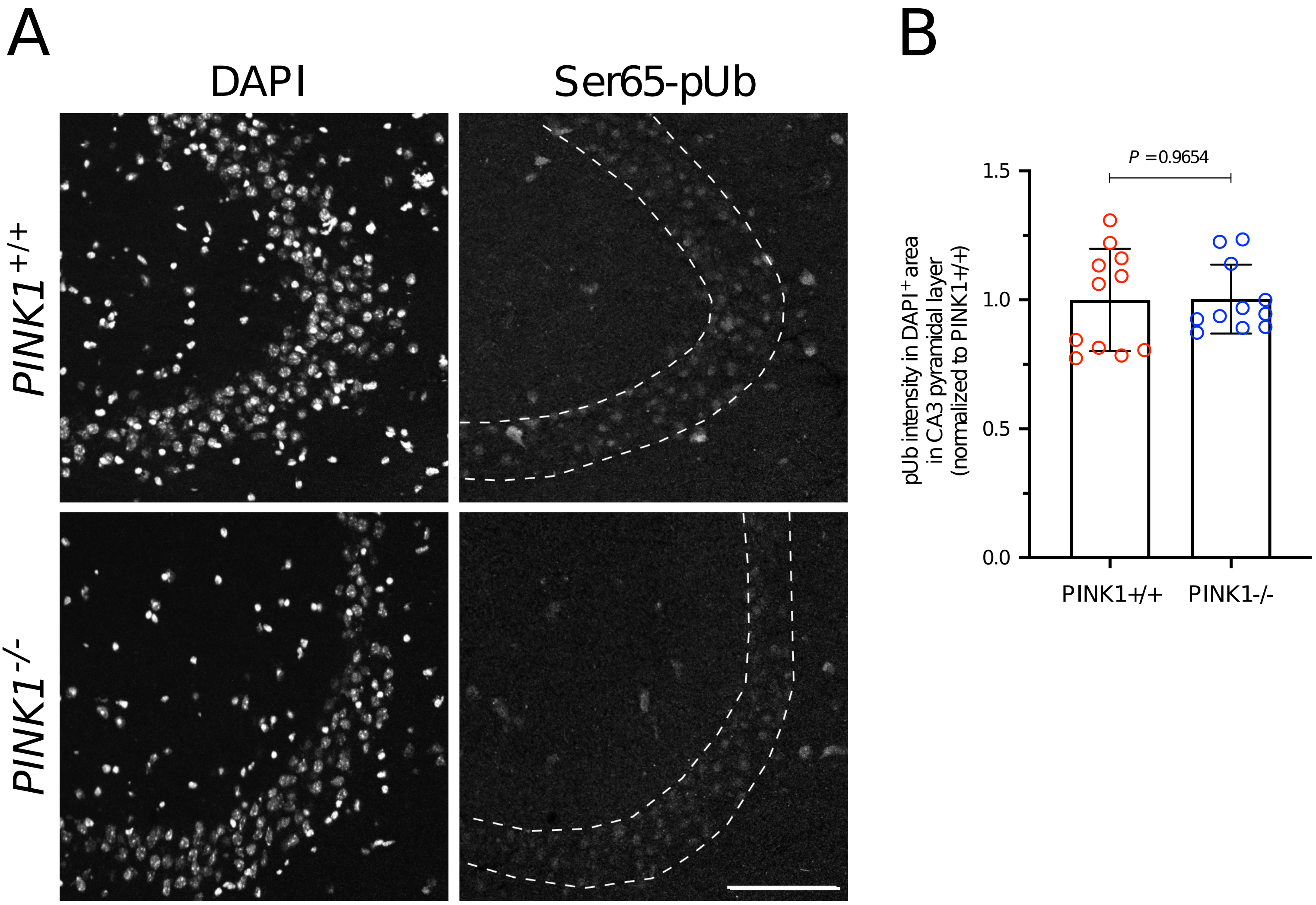

**Supplementary Figure 4. Ser65-pUb signal in DAPI-positive nucleus** (A) Representative confocal microscopy images of hippocampal CA3 regions in PINK1$^{+/+}$ and PINK1$^{-/-}$ mouse. Coronal sections were immunostained with Ser65-pUb antibody. Nucleus was stained with DAPI. Dotted line surrounds CA3 pyramidal cell layer. Scale bar: 100 µm. (B) Quantitative analysis of Ser65-pUb fluorescence signal intensity in DAPI-positive area of CA3 pyramidal cell layer from PINK1$^{+/+}$ and PINK1$^{-/-}$. Data points represent mean signal intensity from one section. Three animals per genotype were used for the analysis.
